# R-loop mapping and characterization during Drosophila embryogenesis reveals developmental plasticity in R-loop signatures

**DOI:** 10.1101/2021.10.29.465954

**Authors:** Alexander Munden, Mary Lauren Benton, John A. Capra, Jared Nordman

## Abstract

R-loops are involved in transcriptional regulation, DNA and histone post-translational modifications, genome replication and genome stability. To what extent R-loop abundance and genome-wide localization is actively regulated during metazoan embryogenesis is unknown. Drosophila embryogenesis provides a powerful system to address these questions due to its well-characterized developmental program, the sudden onset of zygotic transcription and available genome-wide ChIP and transcription data sets. Here, we measure the overall abundance and genome localization of R-loops in early and late-stage embryos relative to Drosophila cultured cells. We demonstrate that absolute R-loop levels change during embryogenesis and that resolution of R-loops is critical for embryonic development. R-loop mapping by strand-specific DRIP-seq reveals that R-loop localization is plastic across development, both in the genes which form R-loops and where they localize relative to gene bodies. Importantly, these changes are not driven by changes in the transcriptional program. Negative GC skew and absolute changes in AT skew are associated with R-loop formation in Drosophila. Furthermore, we demonstrate that while some chromatin binding proteins and histone modification such as H3K27me3 are associated with R-loops throughout development, other chromatin factors associated with R-loops in a developmental specific manner. Our findings highlight the importance and developmental plasticity of R-loops during Drosophila embryogenesis.

## INTRODUCTION

R-loops are a three-stranded nucleic acid structure canonically formed when nascent RNA from transcription reanneals to the template DNA strand, resulting in a displaced single strand of DNA (Aguilera and García-Muse 2012). R-loops were initially identified at the highly transcribed *18S* and *28S* sequences within the rDNA locus of *Drosophila melanogaster* (White and Hogness 1977; Glover and Hogness 1977). More recent studies have demonstrated that R-loops are critical for a diverse set of biological processes (Chédin 2016; Skourtie-Stathaki and Proudfoot 2014). In fact, genome-wide R-loop mapping studies have revealed that R-loops are abundant in eukaryotes and can occupy 10% or more of the genome (Dumelie and Jaffrey 2018; Wahba and Koshland et al. 2016; Fang and Zhang et al. 2019; Xu and Sun et al. 2017; Yan and Liu et al. 2020; Zeller and Gasser et al. 2016; Chen and Fu et al. 2017; Chen and Fazzio et al. 2015; Crossley and Cimprich et al. 2020; Ginno and Chédin et al. 2012; Tan-Wong and Proudfoot et al. 2019; Chan and Hieter et al. 2014; Liu and Han et al. 2021). While R-loops were identified over 40 years ago, their physiological relevance remained elusive for many years.

R-loops are found in all domains of life and their formation is often conserved across cell types and even species (Sanz and Chédin et al. 2016). Deciphering the function of R-loops, however, has been challenging due to their diverse and sometimes contradictory roles in genome function. R-loops are essential for initiation of replication in plasmids and promote mitochondrial genome stability (Dasgupta and Tomizawa et al. 1987; Silva and Aguilera et al. 2018). In contrast, R-loops can block replication fork progression and promote genome instability in an orientation-specific manner (Hamperl and Cimprich et al. 2017; Lang and Merrikh et al. 2017). While potentially causing double-strand breaks at head-on replication-transcription conflicts, R-loops can promote recombination and double strand break repair (Stork and Cimprich et al. 2016; Ouyang and Zou et al. 2021). R-loops also have diverse roles in transcription and chromatin function. In mammalian cells, R-loops have been shown to regulate both histone and DNA methylation at promoter regions (Ginno and Chédin et al. 2012; Chen and Fazzio et al. 2015). While R-loops are often associated with histone modifications correlated with active transcription, recent work has shown that R-loops can help recruit the Polycomb complex to target loci to promote transcriptional silencing (Skourti-Stathaki and Pombo et al. 2019; Alecki and Francis et al. 2020). Genome-wide R-loop mapping studies in yeast, plants and mammalian cultured cells have identified factors such as DNA sequence, DNA topology and histone modifications associated with R-loop formation (Ginno and Chédin et al. 2012; Stolz and Chédin et al. 2019; Hage and Tollervey et al. 2010). R-loop mapping studies in plants and mammalian cells have further revealed that R-loop formation can be dynamic as a function of development (Fang and Zhang et al. 2019; Xu and Sun et al. 2020; Yan and Liu et al. 2020). The extent of R-loop plasticity in other metazoans has yet to be defined. Studying R-loops in the context of development could provide insight into the functional roles R-loops play in establishing developmental-specific changes in chromatin structure, function and transcriptional programs.

Drosophila provide a well-established developmental system to interrogate R-loop plasticity during development. At the earliest stages of Drosophila embryogenesis, rapid cell proliferation is driven by maternally stockpiled proteins and RNA (Tadros and Lipshitz 2009). Approximately two hours after fertilization, zygotic genome activation is triggered and the transcription of over 3000 genes necessary for growth and differentiation are induced in a process known as the maternal-to-zygotic transition (MZT) (Hamm and Harrison 2018; Harrison and Eisen et al. 2011). Prior to the MZT, cells are largely undifferentiated and have abbreviated cell cycles (Foe and Alberts 1983). After the MZT, however, the cell cycle slows and cells become differentiated as morphogenesis proceeds (Farrell and O’Farrell 2014). The changes in cell cycle programs, the onset of zygotic gene activation and cell differentiation during embryogenesis provide a unique system to interrogate whether R-loop formation or resolution impacts embryogenesis and the extent to which, if any, R-loop position and properties change as a function of development.

In this study, we measured R-loop abundance and position in Drosophila embryos and cultured cells. We show that absolute R-loop levels change during embryogenesis and resolution of R-loops is essential for embryogenesis. We mapped R-loops at base pair resolution in 2-3 hour embryos (immediately after the MZT), late-stage embryos (14-16 hours after fertilization) and cultured S2 cells, which are derived from late-stage embryos. We show that, while some sites of R-loop formation are constant during development, there is extensive R-loop plasticity during Drosophila development. Furthermore, we were able to demonstrate changes in the localization of R-loops across gene bodies and the role AT and GC skew play in Drosophila R-loop formation. By leveraging data available through modENCODE and other publicly available datasets, we were able to identify specific histone modifications and chromatin binding proteins associated with R-loop formation in Drosophila and the role active transcription has on R-loop formation. Importantly, developmental-specific R-loops are not driven by transcriptional changes, emphasizing the role that chromatin and R-loop binding proteins play in regulating R-loop formation. Our work establishes Drosophila as a powerful developmental model system to study R-loop biology

## RESULTS

### R-loop abundance is developmentally regulated and R-loop homeostasis is necessary for development

To determine if R-loop abundance and genomic location are regulated throughout development, we turned to the powerful Drosophila embryogenesis system. For our analysis, we chose embryos at two distinct time points: 2-3 hours after egg laying (AEL) and 14-16 hours AEL (Fig. 1A). The 2-3 hour time point corresponds with the onset of the maternal-to-zygotic transition (MZT) occurring during nuclear cleavage cycle 14 (Blythe and Wieschaus 2015). This time point represents the onset of zygotic transcription and allows us to draw upon the wealth of scientific literature that has previously been published, including time-matched modENCODE datasets. The wide-scale activation of zygotic transcription at this time point should provide the first opportunity for R-loop formation during development. To complement this developmental stage, we chose 14-16 hour AEL embryos to understand how R-loop formation might differ in differentiated cells with a more mature chromatin environment and a transcription program characterized by cell-type-specific maintenance (Bonnet and Müller et al. 2019; Bowman and Bender 2014; Smith and Orr-Weaver 1991). S2 cells, an established Drosophila cell culture line derived from late-stage embryos (Schneider 1972), were used to determine how R-loops might differ between embryos and cultured cells, where the majority of R-loop research has been conducted.

**Figure 1:**
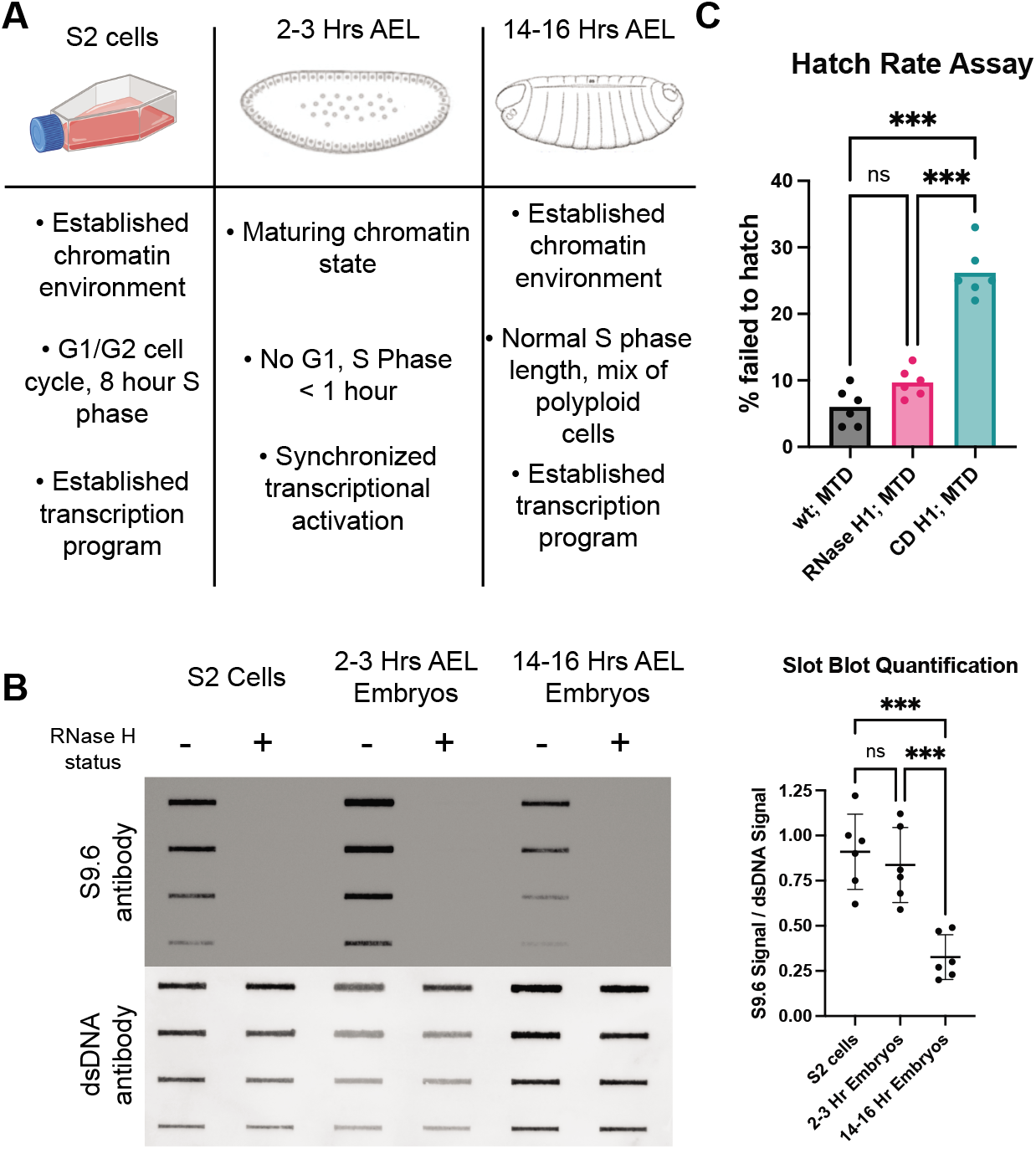
R-loop abundance is developmentally regulated and R-loop homeostasis is necessary for development. (A) Schematic summarizing how the chromatin environment, developmental stage, and replication program vary among the developmental samples used. (B) Representative slot blot of RNA:DNA hybrid levels, measured by S9.6 antibody intensity, across samples. RNase H treatment verifies specificity of antibody, and antibody specific for double-stranded DNA is used as a loading control. Quantification of signal for six biological replicates is to the right. *** < 0.05, one-way ANOVA with Tukey’s multiple comparisons test. (C) Hatch rate among embryos that overexpress RNase H1 (H1) or a catalytic dead RNase H1 (CD). *** < 0.05, one-way ANOVA with Tukey’s multiple comparisons test.

To begin, we asked whether the absolute levels of R-loops are influenced by development. To this end, genomic DNA was extracted from each sample and spotted onto a nitrocellulose membrane and probed with the S9.6 antibody, which recognizes RNA:DNA hybrids (Boguslawski and Carrico 1986). S2 cells and 2-3h embryos showed similar amounts of S9.6 signal, while DNA from 14-16h embryos showed a significant decrease in S9.6 signal (Fig. 1B). To ensure that the S9.6 signal stems from R-loops, we pretreated control samples with RNase H, which degrades the RNA moiety of a RNA:DNA hybrid. The S9.6 antibody has some specificity to double-stranded RNA and Drosophila embryos are known to contain dsRNA (Hartono and Vanoosthuyse et al. 2018). In fact, in the RNase H treated control samples we initially detected some signal with the S9.6 antibody, which was completely eliminated by pretreatment with RNase III. Therefore, for all R-loop assays we pretreat our samples with RNase III to ensure S9.6 signal isn’t due to dsRNA.

Next, we asked whether perturbing R-loop homeostasis affects embryogenesis. To answer this, we generated flies that overexpress a GFP-tagged, nuclear localized version of Drosophila RNase H1 or a catalytically dead version of the same protein (RNase H1^CD^). To ensure that the RNase H1 proteins were maternally deposited and present at the earliest stages of embryogenesis, we used the pUASz expression system coupled with the maternal triple driver (DeLuca and Spradling 2018; Rørth 1998). After confirming that the GFP was observable by western blot (Supplemental Fig. 1), we performed a hatch rate assay to determine if perturbing R-loop homeostasis affects embryogenesis. We observed a consistent but statistically insignificant hatching defect in the RNase H1 overexpression embryos (Fig. 1C). The RNase H1^CD^ expressing embryos, however, had a ~25% failure to hatch rate, which was significantly different from the wild-type and the RNase H1 overexpression controls. Overall, we conclude that the absolute abundance of R-loops changes during development and that preventing R-loop processing through overexpression of a catalytically dead RNase H1 results in embryonic lethality.

### R-loop position and properties are influenced during development

While the absolute abundance of R-loops changes during development, we wanted to determine how R-loop position throughout the genome changes during Drosophila development. Genome-wide R-loop mapping during Drosophila development would allow us to ask if R-loop formation is hardwired into the genome driven only by cell-type-specific transcription, or, more interestingly, is R-loop formation plastic during development changing independent of sequence composition and transcription status. To address this question, we performed <u>D</u>NA:<u>R</u>NA <u>i</u>mmuno<u>p</u>recipitation on sonicated nucleic acids followed by strand-specific sequencing of the DNA strand (ssDRIP-seq) in S2 cells, 2-3h and 14-16h embryos (Fig. 2A) (Xu and Sun 2017). We initially tried DNA-RNA immunoprecipitation followed by cDNA conversion coupled to high-throughput sequencing (DRIPc-seq) (Sanz and Chédin et al. 2016). When conducted in Drosophila, however, we found high levels of RNA contamination in the final sequencing results (data not shown). Even with the ssDRIP-seq method, it was necessary to pre-treat genomic DNA preps with RNase A and RNase III as Drosophila embryos are stockpiled with RNA.

**Figure 2:**
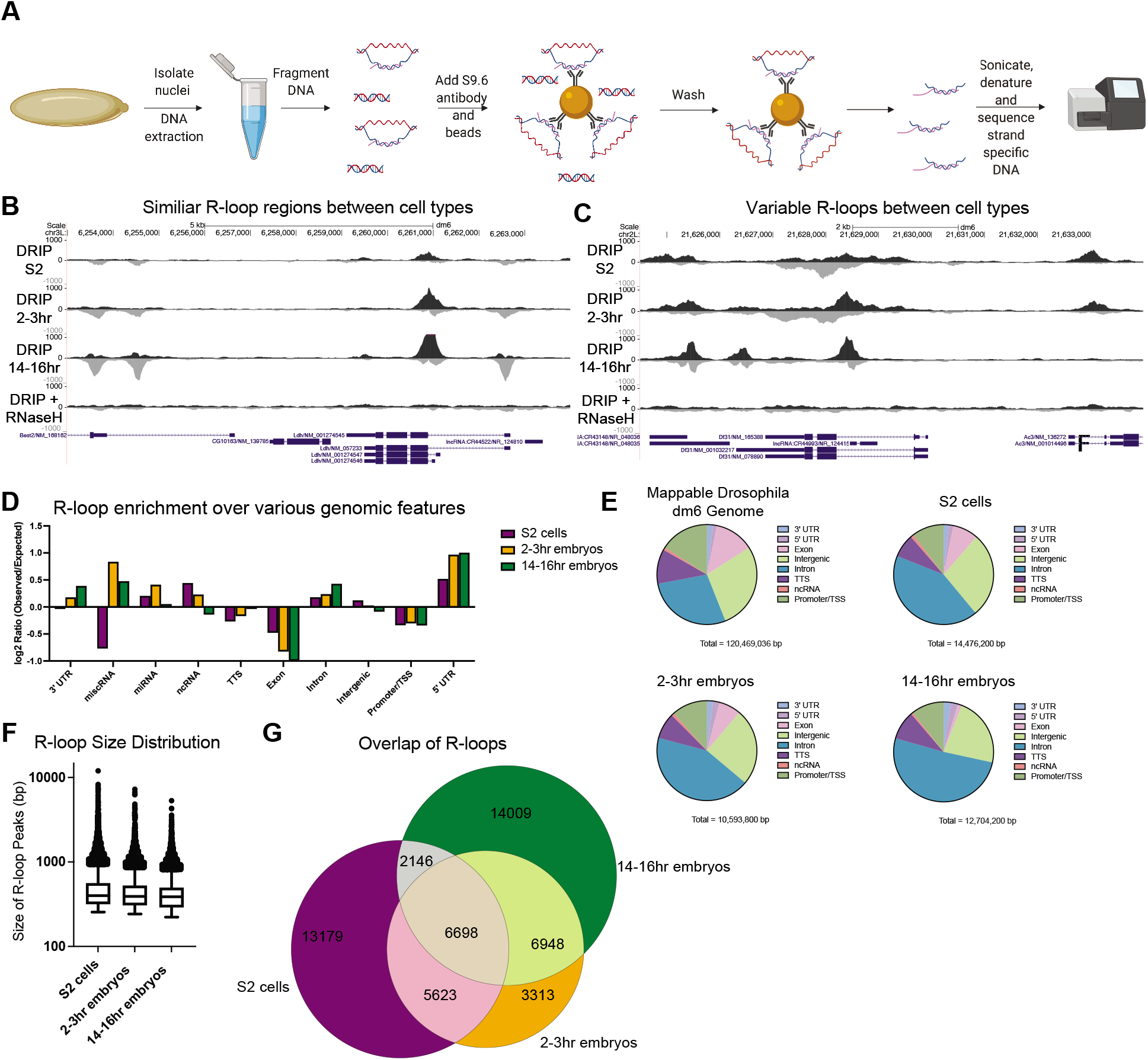
The R-loop landscape changes as a function of development. (A) Diagram of the ssDRIP-seq mapping strategy. (B) ssDRIP-seq snapshot of a 10kb region on chromosome *3L* where R-loop distribution is similar between samples. (C) ssDRIP-seq snapshot of a 10kb region on chromosome *2L* where R-loop distribution varies between samples. (D) R-loop enrichment relative to the expected distribution for common genomic features. (E) R-loop abundance within indicated genomic regions for each developmental sample. (F) The distribution of R-loop sizes at different timepoints for each developmental sample. (G) Overlap of R-loops between developmental samples.

ssDRIP-seq of embryos and S2 cells revealed strand-specific signal that was sensitive to RNase H pretreatment, and showed cell-type specific R-loop formation (Fig. 2B and 2C). Biological replicates were highly correlated (Supplemental Fig. 2A) and our ssDRIP data sets were well correlated with recently published ssDRIP-seq data sets in Drosophila S2 cells and embryos, although different time points were used (2-3h and 14-16h vs. 2-6h and 10-14h embryos) (Alecki and Francis et al. 2020). We validated several sites using DRIP-qPCR to confirm our sequencing results (Supplemental Fig. 2B). These data indicate that our ssDRIP signal reflects RNA:DNA hybrid position throughout the genome and ssDRIP is a robust method to map sites of R-loop formation in Drosophila.

To map the precise location of R-loops throughout the genome and allow us to compare both quantitative and qualitative properties of R-loops, we used MACS to define R-loop peaks (Zhang and Lui et al. 2008). Peaks were called separately against the input samples and RNase H treated controls, and all overlapping peaks were kept for analysis. Using this criterion, we identified 28,464, 22,581 and 28,961 peaks in S2 cells, 2-3h and 14-16h, respectively, which occupied between 8.3 and 12.5% of the genome. R-loop peak size was similar between sample types with a median of approximately 500 bp, but R-loops could occupy zones up to 10kb in size (Fig. 2F). Out of the 51,916 total unique R-loop peaks identified between all samples, 12.9% were common to all sample types, 28.3% were present in at least two samples and 58.8% were specific to an individual sample (Fig. 2G).

Since ssDRIP allows for strand-specific annotation, we characterized R-loops relative to strand-specific genomic features. Relative to transcription units, ~35% of R-loops occur in sense to transcription in S2 cells and 2-3h embryos, whereas ~30% of R-loops are antisense (Supplemental Fig. 2C). Interestingly, in the 14-16h embryos, a greater fraction of R-loops occurs antisense relative to transcription (~40%; Supplemental Fig. 2C). In all samples, 30-35% of the R-loops form in unannotated regions of the genome. Next, we used HOMER to annotate R-loop signal relative to genomic features (Heinz and Glass et al. 2010). In all samples, we found that R-loops are enriched in the 5’ UTR, introns and in miRNA regions, while R-loops are universally depleted in exonic regions (Fig. 2D-E). The depletion of R-loops in exons provides additional support that our R-loop peaks are not an artifact of RNA contamination (Fig. 2D and 2E). Consistent with previous R-loop mapping studies, we identified strong R-loop signal at the rDNA locus and the histone gene locus (Supplemental Fig. 2D and 2E) (Constantino and Koshland 2018; Dumelie and Jaffrey 2017). We also observed developmental-specific differences in R-loop formation. For example, R-loop signal was enriched in miRNA and ncRNA regions only in S2 cells and 2-3h embryos. Taken together, these results demonstrate that R-loop signal across Drosophila development is dynamic.

### R-loop enrichment at transcription units changes during development

In mammals, R-loops are known to preferentially form at transcription start sites (TSS), gene bodies and transcription termination sites (TTS) (Sanz and Chédin et al. 2016; Skourti-Stathaki and Proudfoot et al. 2014). To ask if this pattern of R-loop formation is similar in Drosophila, and whether it changes during development, we measured R-loop abundance across gene bodies in our developmental samples. S2 cells and 2-3h embryos display a very similar pattern of R-loop formation, with a strong peak at the TSS and continued signal over the gene body (Fig. 3A), which is similar to R-loop positions in other metazoans (Sanz and Chédin et al. 2016). Interestingly, there is a depletion of R-loops immediately after the TTS in S2 cells (Fig. 3A). The 14-16h embryos, however, have a significantly different pattern altogether, with R-loop enrichment at the TSS, lower signal over the gene body relative to S2 cells and 2-3h embryos and a strong enrichment at the TTS (Fig. 3A). To determine if these patterns were driven by sense or antisense R-loops, we generated metaplots using strand-specific data. This analysis revealed that sense R-loops recapitulate this pattern, except with the 2-3h embryos having a more pronounced signal over the gene body. In general, antisense R-loops have a stronger signal at the TTS. In the 14-16h embryos, however, the majority of the signal at the TSS and TTS is derived from antisense R-loops (Fig. 3A). Taken together, we conclude that R-loop enrichment at transcription units is not hardwired into the genome, but can be dynamic as a function of development.

**Figure 3:**
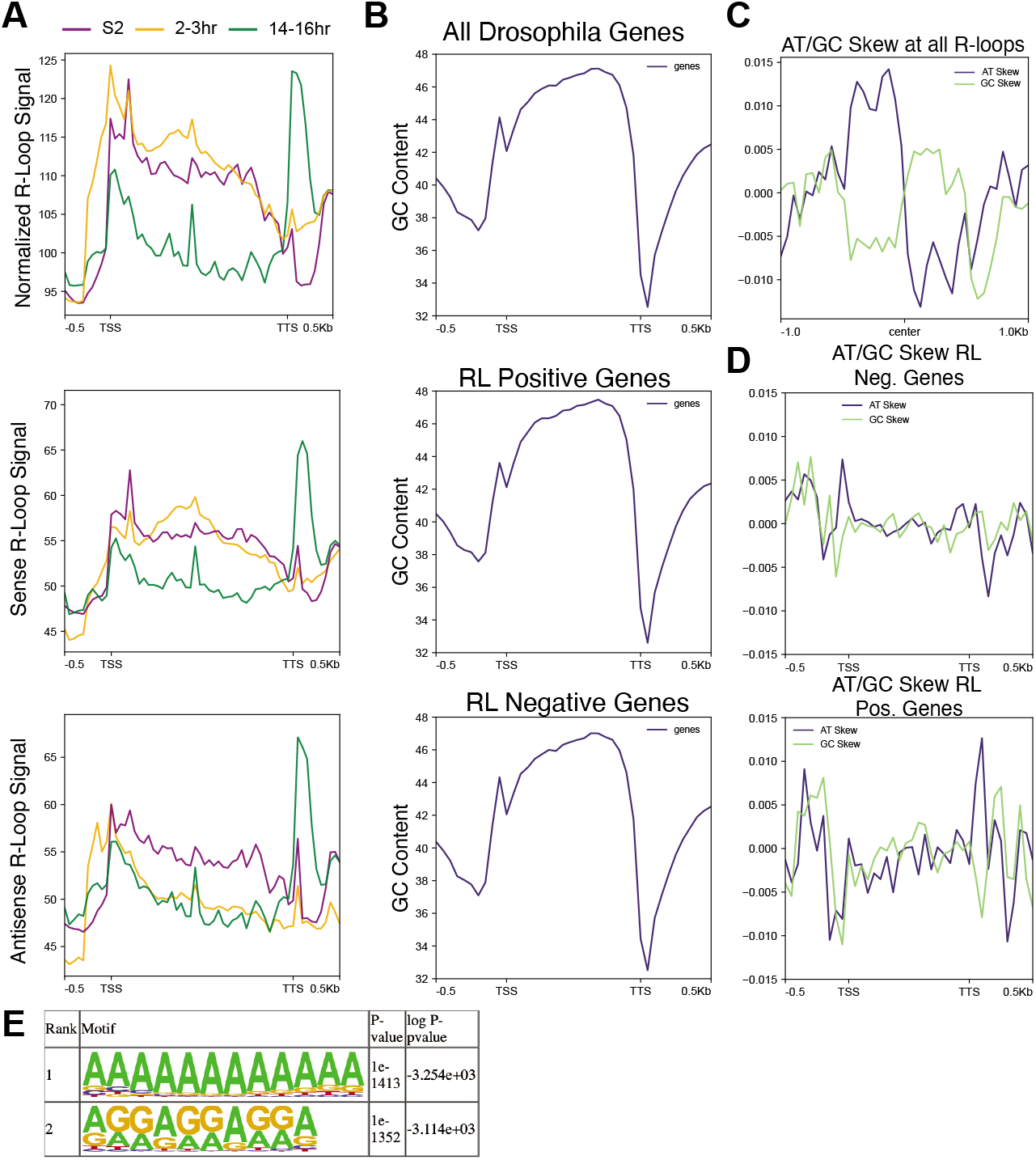
R-loop signal as a function of transcription unit and sequence composition. (A) Metaplot of ssDRIP-seq signal for all samples relative to the gene body. Top panel is total R-loop signal, middle panel is sense R-loops, bottom panel is anti-sense R-loops. (B) The GC composition of all Drosophila genes, genes that have an R-loop in one of the developmental samples and genes that lack any R-loop signal. (C) Metaplot of GC and AT skew across all identified R-loops. (D) Metaplot of GC and AT skew across the gene body of genes that lack R-loops (top) and genes that form an R-loop. (E) DNA sequence motifs in the peaks of all R-loops identified my HOMER.

Given that the absolute levels and relative position of R-loops can change between developmental states in Drosophila, we wanted to assess the contribution DNA sequence composition has on R-loop formation in Drosophila. Unlike in mouse and human cells, Drosophila lack high GC content at the TSS. In fact, GC content decreases relative to the gene body in Drosophila (Fig. 3B). We asked if R-loop forming genes differ in their GC content relative to genes that lack R-loops. We found that genes with and without R-loops have a near-identical GC content along the gene body (Fig. 3B). While overall GC content is not different in R-loop positive or negative genes, GC and AT skew has been shown to be a contributing factor to R-loop formation (Ginno and Chédin et al. 2012). To test if GC or AT skew is associated with R-loop formation in Drosophila, we measured the AT/GC skew directly over all identified R-loops. This analysis revealed a striking transition from positive to negative AT skew at the center of our combined R-loop signal. This is mirrored by a transition from negative to positive GC skew centered at the combined R-loop signal (Fig. 3C). Highlighting the robustness of this transition in skew, even developmental-specific R-loops display the same transition in AT/GC skew (Supplemental Fig. 3A).

We also calculated GC and AT skew for R-loop forming and deficient genes in all samples. Stronger negative GC skew at the TSS and TTS were observed in R-loop forming genes relative to genes that fail to form R-loops (Fig. 3D). Specifically, AT skew at the TSS transitioned from positive skew in R-loop deficient genes to negatively skew in R-loop forming genes. At the TTS, there is a strong positive AT skew immediately downstream of the TTS only in R-loop forming genes (Fig. 3D). Negative GC skew is stronger in at both the TSS and TTS in R-loop forming genes. This analysis reveals a correlation between altered AT skew and negative GC skew in R-loop forming genes, suggesting that AT/GC skew could contribute to R-loop formation in Drosophila. Together, we conclude that while AT and GC skew could facilitate R-loop formation, developmental-specific R-loop formation is not likely driven by changes in AT or GC skew. This suggests that transcription, chromatin environment or other factors could contribute to cell type specific R-loop formation.

To test whether any specific DNA sequence motifs are associated with R-loop formation, we searched for motifs enriched in the set of all Drosophila R-loops. Two motifs stood out as an order of magnitude more significantly enriched that any others: a polyadenine tract and a polypurine tract (Fig. 3E, Supplemental Fig. 3B for the entire table). This indicates that polypurine tracts are conducive to R-loop formation, which is consistent with the known thermodynamic stability of RNA:DNA hybrid formation in purine-rich template sequences (Huppert 2008).

### Common and cell-type specific chromatin features associated with R-loops

R-loops are associated with activating chromatin marks such as H3K4me1/2/3 and H3K9ac and, to a lesser extent, with repressive chromatin marks such as H3K27me3 (Sanz and Chédin et al. 2016). Chromatin marks associated with R-loops, however, vary depending on species. One possibility is that there are marks that are universally associated with R-loop formation whereas some chromatin marks could associate with R-loops in a developmental-specific manner. To answer this question, we leveraged time-matched ChIP-seq modENCODE datasets for S2 cells, 2-4h embryos (ChIP-chip and ChIP-seq) and 14-16h embryos. To quantitatively determine if chromatin marks were positively or negatively associated with R-loops, we evaluated the probability of R-loops overlapping a variety of histone modifications and chromatin-associated proteins by chance using a peak shuffling bootstrap procedure (see Materials and Methods). The available chromatin proteins vary for each sample, but there are 10 chromatin or histone markers common in all three developmental samples (Fig. 4A). Several factors that are associated with transcriptional activation, and have been previously shown to be associated with R-loops, are enriched at R-loops in S2 cells and 2-3 hour embryos (Fig. 4A, Supplemental Fig. 4). Additionally, repressive chromatin marks such as Polycomb complex subunits and H3K27me3 are enriched in all samples, which is consistent with recent work linking R-loops to transcriptional repression (Fig. 4A, Supplemental Fig. 4) (Skourti-Stathaki and Pombo et al. 2019; Alecki and Francis et al. 2020).

**Figure 4:**
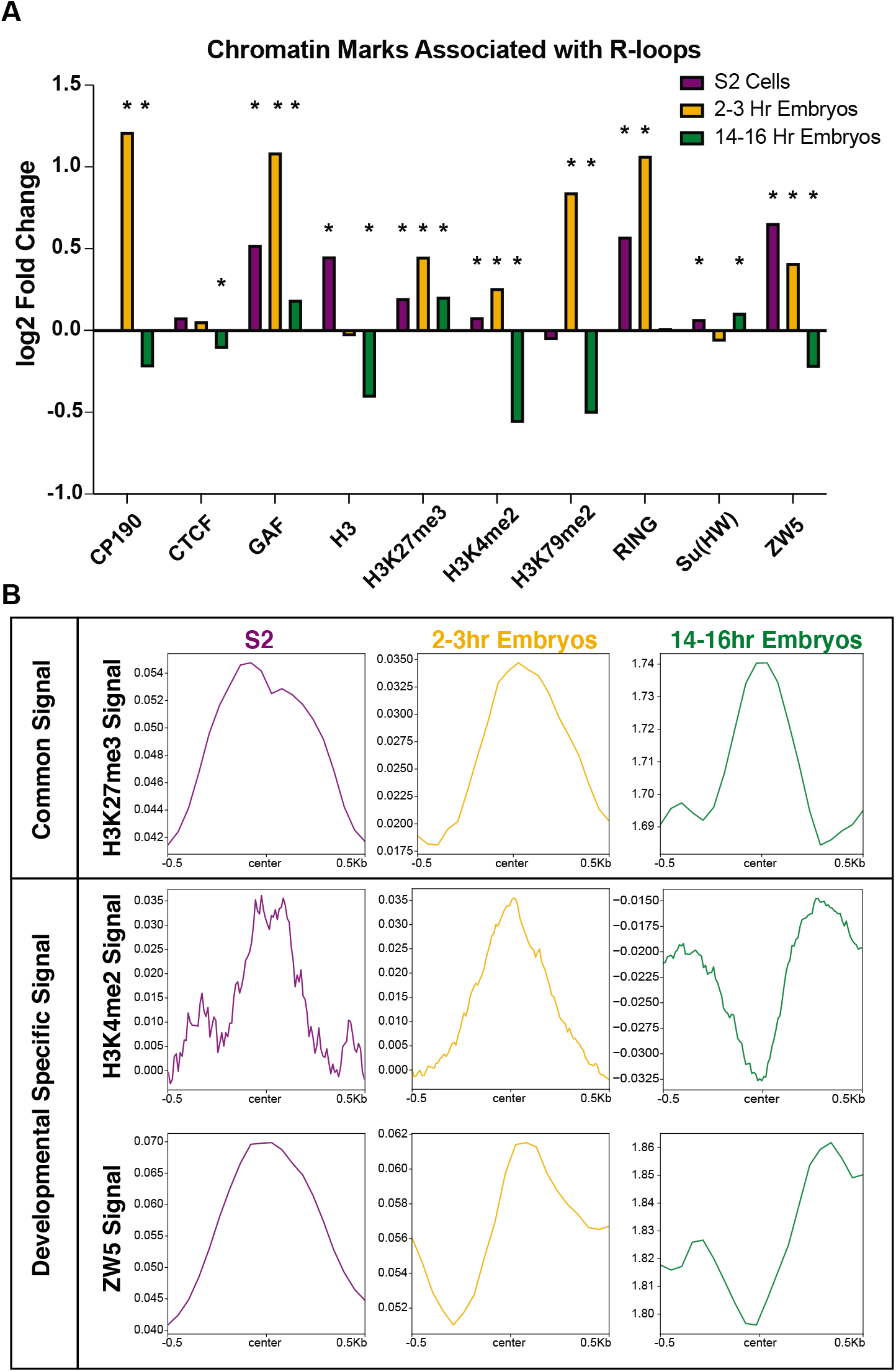
Common chromatin features associated with R-loops. (A) Log2 fold enrichments of chromatin-associated factors within R-loop regions in common for S2 cells, 2-3 hour embryos and 14-16 hour embryos. * < 0.05 with Bonferroni correction for multiple testing (B) Metaplots of H3K27me3, H3K4me2, and ZW5 ChIP-chip (S2 and 2-4 hour embryos) and ChIP-seq (14-16 hour embryos) confirming common and developmental-specific enrichment of chromatin factors at R-loops.

We asked which marks are consistently associated with R-loops (positively or negatively) across development and which factors are developmental specific. We found that the repressive mark H3K27me3 was positively associated with R-loops in all developmental samples, highlighting the link between R-loops and transcriptional repression (Fig. 4B). Interestingly, we identified factors (H3K4me2 and ZW5) that were enriched in one developmental sample but not in others (Fig. 4B). These results suggest while some factors are associated with R-loops regardless of development state, other factors are associated with R-loops in a developmentally-specific manner.

### R-loop formation as a function of transcription

In this study, we have noted distinctive changes in R-loop formation across development. Once possibility is that these changes are driven by developmental-specific changes in the transcription program. As embryos are stockpiled with maternally deposited RNA and RNA-seq is an indirect readout of active transcription, we turned to previously published and time-matched GRO-seq datasets in S2 cells and 2-2.5h embryos, respectively (Core and Lis et al. 2012; Saunders and Ashe et al. 2013). Unfortunately, time-matched GRO or PRO-seq datasets do not exist for 14-16h embryos. We converted GRO-seq signal to FPKM for each annotated transcript in the Drosophila transcriptome. Then, we compared the GRO-seq value of all R-loop-containing genes to genes devoid of R-loops. In S2 cells, R-loop positive and negative genes had a similar median FPKM value by GRO-seq (Fig. 5A). R-loop-containing genes in 2-3h embryos, however, revealed a different paradigm. R-loop positive genes had a significantly higher expression level than R-loop negative genes (Fig. 5C). To ask if R-loop-containing genes were over or underrepresented with genes that have high or low expression levels, we binned GRO-seq FPKM values into quartiles and asked what fraction of R-loop containing genes fell within each expression quartile (Fig. 5B, D). In S2 cells, R-loop containing genes were slightly overrepresented in the highest expression quartile and, to a lesser extent, in the lowest expression quartile (Fig. 5B). In 2-3h embryos, however, R-loops were significantly overrepresented in the highest expression quartile and underrepresented from the lowest expression quartile (Fig. 5D). While analyzing this data, we also found the number of R-loops forming sites per gene was correlated with transcriptional activity (Fig. 5E). We observe a consistent increase in the average number of R-loops per gene as transcriptional activity increases (Fig. 5E). The increase in the average number of R-loops per gene could represent multiple R-loops within a given gene or larger R-loop zones allowing R-loops to form over a larger target region.

**Figure 5:**
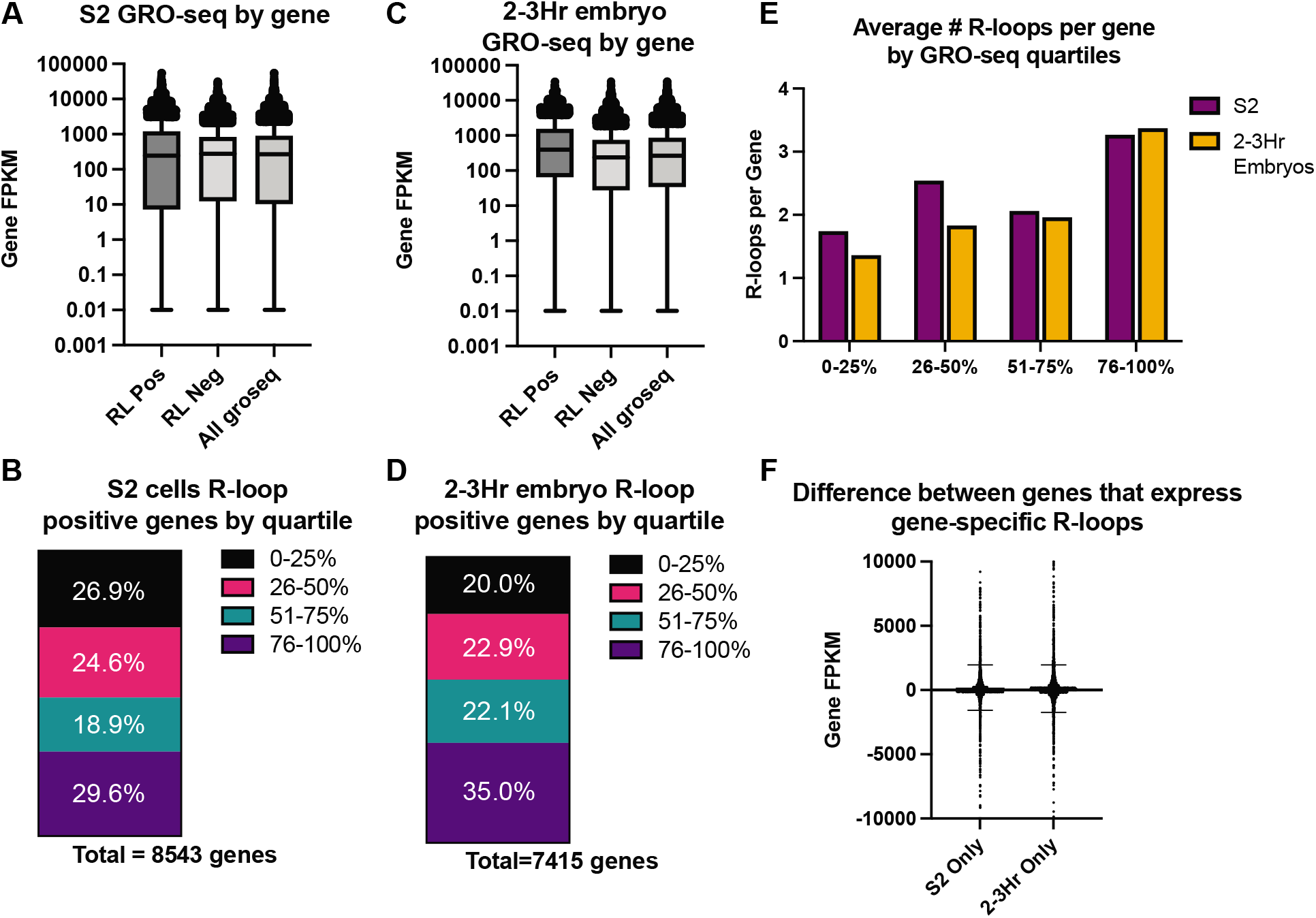
R-loop formation as a function of transcription. (A) GRO-seq values for genes that contain R-loops (RL Pos) and genes that do not contain R-loops (RL Neg) in S2 cells. (B) Transcripts were sorted into quartiles based upon GRO-seq expression, and R-loop forming genes were assigned to their respective quartile. (C) Same as A, except for 2-3 hour embryos. (D) Same as B, except for 2-3 hour embryos. (E) The average number of R-loops detected for each gene in each of the expression quartiles is graphed for S2 cells and 2-3 hour embryos. (F) The difference in GRO-seq values between S2 cell and 2-3 hour embryos were queried for genes that showed developmental-specific R-loop formation.

One explanation for developmental-specific R-loop formation is that specificity is driven by developmental-specific transcription status. To test this, we compared expression level of genes that exhibit R-loops only in S2 cell or only in 2-3h embryos (Fig. 5F). If active transcription drives the changes in R-loop formation, we would expect R-loop positive genes that are unique to 2-3h embryos would have significantly higher expression level in 2-3h embryos relative to S2 cells, and vice-versa. The median difference of GRO-seq values in developmental-specific R-loop-containing genes, however, is approximately zero with a normal distribution (Fig. 5F). Therefore, we conclude that active transcription is not a driver of developmental-specific R-loop formation and that factors such as chromatin state or R-loop-specific proteins drive these differences.

### R-loops have the potential to trigger ATR activation at the MZT

The onset of zygotic transcription at the MZT is associated with RPA accumulation at the 5’ end of genes and activation of the ATR-mediated DNA damage checkpoint response (Blythe and Wieschaus, 2015). Delaying the onset of zygotic transcription delays the activation of ATR (Mei41 in Drosophila), indicating that replication-transcription conflicts drive the activation of the DNA damage response that occurs at the MZT (Blythe and Wieschaus, 2015; Sibon and Theurkauf et al. 1999). It is unknown, instability at the MZT, we would predict to see an enrichment of RPA at R-loop forming sequences in 2-3h embryos. Qualitatively, we see overlap between RPA and R-loops in 2-3h embryos (Fig. 6A). We tested the significance of this overlap by using the random shuffling method previously described. Quantitatively, we observe a significant enrichment of RPA at R-loop forming sequences in the 2-3h embryo. Importantly, there was an even more substantial enrichment of RPA at R-loop peaks that are unique to 2-3h embryos (Fig. 6B). This data suggests that R-loops could contribute to the transcription-induced DNA damage that occurs in the absence of ATR at the MZT. We do note, however, that the RPA ChIP-seq data comes from a time point ~20 minutes earlier in development than the time point we chose for R-loop mapping (Blythe and Wieschaus, 2015). Given this caveat, we think it is even more notable that significant overlap of RPA and R-loops is observed in this analysis.

**Figure 6:**
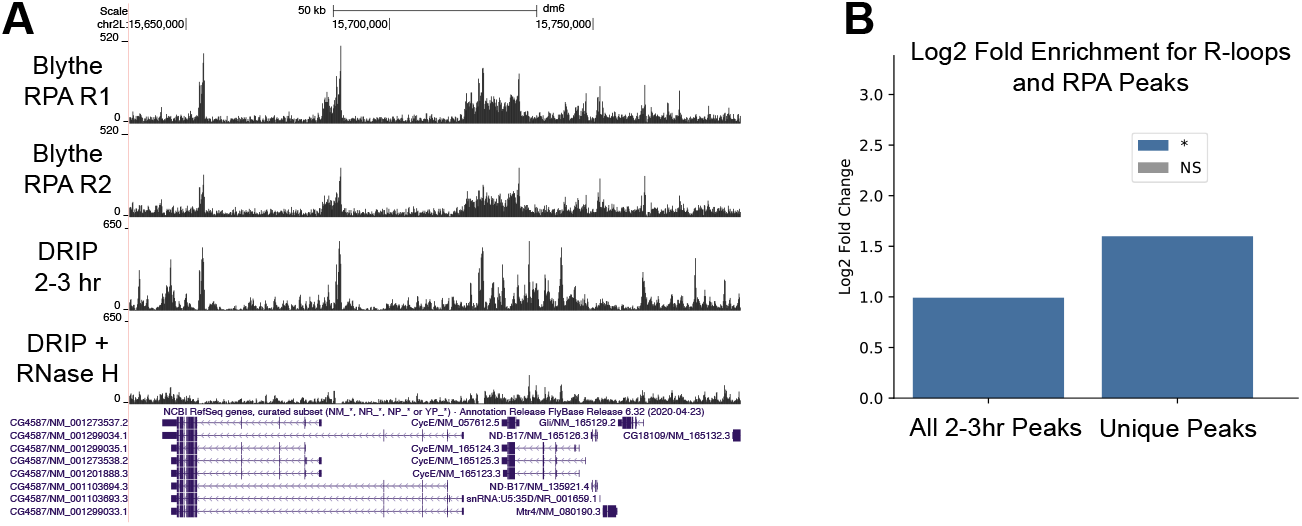
R-loops have the potential to trigger ATR activation at the MZT. (A) Overlap of RPA ChIP-seq profiles from cycle 13 embryos (Blythe and Wieschaus et al. 2015) and ssDRIP-seq profiles from 2-3h embryos. (B) Log2-fold enrichment of RPA at all 2-3h R-loop peaks or R-loops that are specific for 2-3h embryos.

## DISCUSSION

By mapping R-loops in a developing organism, we have been able to provide new insight into the role that DNA sequence, active transcription and chromatin associated factors has on R-loop formation. While previous analysis of R-loop metabolism across development has been performed in plants and mammalian cultured cells (Yan and Liu et al. 2020; Xiu and Sun et al. 2020; Shafiq and Sun et al. 2017), we present the first functional characterization of R-loops during Drosophila embryogenesis. A developmental approach to studying R-loop formation is that it allows the distinction between factors that are stably linked to R-loop formation from those that are developmental specific. This has the potential to identify key molecules and processes that could drive R-loop formation and resolution during development and disease.

One surprising finding is that the absolute level of R-loops changes during embryogenesis. This is unlikely due to changes in transcription during development as the stages of embryogenesis used in this study are similarly active. This suggests that there is an active mechanism which prevents R-loop formation or resolves active R-loops during later stages of Drosophila embryogenesis. The importance of R-loop processing during development is further highlighted by the observation that preventing R-loop degradation by overexpression of a catalytically inactive version of RNase H1 causes hatching defects in Drosophila embryos. Interestingly, overexpression of catalytically active RNaseH1 did not have the same effect. One possible explanation of this result is that hyper stable R-loops block replication, causing genome instability (Stork and Cimprich et al. 2016; Lang and Merrikh et al. 2017). Alternatively, hyper stable R-loops could drive chromatin or transcriptional changes that negatively impact embryogenesis (Lima and Crooke et al. 2016). Further work will be required to distinguish between these and other possibilities.

Specific DNA sequence biases are associated with R-loop formation (Ginno and Chédin et al. 2012; Stolz and Chédin et al. 2019). While we found that overall GC content is the same for R-loop positive and negative genes, AT and GC skew were associated with R-loop forming sequences. Interestingly, this skew varied as a function of the transcription unit. Promoter-associated R-loops have low AT and GC skew, whereas R-loops in transcriptional termination regions have high AT skew, but low GC skew. This was unexpected given that G4 quadraplex forming regions with high GC skew on the non-template strand are associated with R-loop formation (Ginno and Chédin et al. 2012; Lee and Myong et al. 2020). Additionally, R-loops can modulate DNA methylation at CpG islands in promoter regions (Ginno and Chédin et al. 2012). Unlike in plants and mammals, however, Drosophila lack wide-scale DNA methylation (Capuano and Ralser et al. 2014). Therefore, Drosophila allows the uncoupling between R-loop formation and DNA methylation, which could explain why R-loops are associated with a higher AT skew than GC skew in Drosophila. Similar to other organisms, however, we have found several polypurine motifs associated with R-loops. This likely reflects the thermodynamic stability associated with RNA:DNA hybrids at purine-rich sequences (Huppert 2008). AT and GC skew can also vary as a function of a transcription unit, with promoter regions having higher GC skew that the gene body or termination region. One interesting observation in Drosophila is that the R-loop signal relative to the transcription unit can vary as a function of development. The most significant difference is in 14-16h embryos where R-loops are enriched at TTS, but not in 2-3h embryos or S2 cells. This difference does not appear to be driven by AT or GC skew. We propose that a combination of factors such as transcription status, chromatin marks and R-loop binding proteins drive these changes in R-loop formation during development.

We have found that R-loops are positively and negatively associated with specific histone modifications and chromatin associated factors. Many of the factors we analyzed in Drosophila have been shown to be enriched or depleted in other systems, including mammalian cells (Sanz and Chédin et al. 2016; Pinter and Rathert et al. 2021; Herrera-Moyano and Aguilera et al. 2014). More importantly, however, factors associated with R-loops can change as a function of development. For example, R-loops in 14-16h embryos lose their association with common activating histone marks such as H3K4me3 and H3K36me2/3. In contrast, H3K27me3 is enriched at R-loops in all developmental states. Therefore, it is critical to assay multiple cell types or developmental states before concluding that a chromatin factor is correlated with R-loop formation.

The link between R-loops, transcription state, histone marks and chromatin associated factors has been seen in other organisms (Sanz and Chédin et al. 2016). In Drosophila, we see a consistent relationship between active and repressive chromatin marks, signified by enrichment in both H3K27ac and H3K27me3, and R-loop formation. This is supported by the association of R-loops with both highly active and silent genes in both embryos and cultured cells. Our work, and that of others, identify R-loops associated with transcriptionally active and inactive genes (Skourti-Stathaki and Pombo et al. 2019). This suggests that, at least in Drosophila, there may exist at least two classes of R-loops. R-loops that form as a byproduct of active transcription and R-loops that function in a repressive capacity to prevent transcription within repressive chromatin domains. This would be consistent with recent work demonstrating that R-loops facilitate silencing by the Polycomb complex (Alecki and Francis et al. 2020; Skourti-Stathaki and Pombo et al. 2019). Additionally, the abundance of R-loops in LTR and LINE elements in early embryos support the idea that R-loops prevent transcription of these elements (Zeller and Gasser et al. 2016; Bayona-Feliu and Azorín et al. 2017; Zeng and Hamada et al. 2021). Understanding how different categories of R-loops maintain their identity will be an exciting challenge. For example, how do cells know which R-loops should function in a repressive manner versus those that function as activators? The question of whether R-loops help establish a chromatin state or are a function of it remains an outstanding question in R-loop biology.

Mapping of R-loops has been performed in a variety of organisms ranging from yeast, worms, plants, and mammalian cultured cells. While there are factors and processes that are consistently associated with R-loops across organisms, there are also key differences. For example, in plants there are low levels of R-loops at gene terminators compared to other organisms and high accumulation of antisense R-loops that regulate specific loci (Xu and Sun et al. 2020; Sun and Dean et al. 2013). In contrast, mammalian cells exhibit R-loops at promoters and TTS and the number of antisense R-loops are much more limited (Sanz and Chédin et al. 2016). The fact that Drosophila exhibit changes in antisense R-loop signal across the gene body depending on developmental state highlights the importance of examining R-loops in a developmental context. Drosophila provides a powerful model to understand key properties of R-loop biology in the context of unperturbed metazoan development. Here, we demonstrate that R-loop formation within the same genomic sequence can vary as a function of development. Our work suggests that a combination of transcription, chromatin-associated factors and sequence elements drive differential R-loop formation during development. Therefore, Drosophila provides a powerful model to understand, mechanistically, the factors responsible for R-loop formation and resolution to execute specific developmental programs.

## METHODS

### S9.6 antibody

A hybridoma cell line producing the S9.6 antibody was purchased through ATCC (product #HB-8730). The cell line was grown under recommended conditions. The S9.6 antibody was purified on a protein G column using the GE aKTA system and run over a desalting column for buffer exchange into PBS to obtain a final concentration of 1 mg/mL. The antibody was aliquoted and stored at −80°C. A fresh aliquot was used for every ssDRIP-seq experiment.

### RNase H1 overexpression

Drosophila RNase H1 was cloned from RNA derived from Oregon R embryos. RNA was converted into cDNA, PCR amplified, and cloned into the pUASz vector with a C-terminal GFP tag (DeLuca and Spradling 2018). The A isoform was chosen as the isoform B isn’t detected in Drosophila tissues (Cózar de and Jõers et al. 2019). The mitochondrial localization start site was converted to AAA to ensure RNase H1-GFP would only be present in the nucleus. The catalytically dead version of RNase H1 (D201N) was made by site-directed mutagenesis (Agilent QuickChange Lightning). Plasmids were injected into an *attP2* containing stock (BestGene) for site-specific integration.

### Hatch rate assay

For the overexpression experiments, homozygous RNase H1 males were crossed with unmated female homozygous for the maternal triple driver (MTD, Bloomington Stock 31777) to drive expression early in embryogenesis. Male Oregon R flies were crossed with MTD females as a control. Progeny were transferred to bottles with a grape juice agar plate with wet yeast for embryo collection. 100 unhatched embryos were carefully moved to a fresh grape juice plate and incubated overnight at 25°C. After 36h, unhatched embryos were counted. This was repeated three times each from two separate crosses.

### Cell culture

S2 cells were obtained directly from the Drosophila Genomic Resource Center (DGRC). Cells were confirmed negative for mycoplasma contamination via PCR. Cells were grown at 25°C in Schneider’s Drosophila Medium with 10% heat-inactivated FBS (Gemini Bio Products) and 100 U/mL of Penicillin/Streptomycin (Fisher Scientific).

### Embryo collection and staging

Oregon R flies were expanded into population cages containing grape juice plates supplemented with wet yeast. Population cages were kept at 25°C in a humidified room and plates were changed daily. Before embryo collections, flies were precleared for at least one hour to minimize the number of late-stage embryos. Embryos were collected and aged at 25°C to obtain embryos that were 2-3 or 14-16 hours old. After aging and collection, embryos were dechorionated in 50% bleach for 2 minutes and thoroughly rinsed in water. Embryos were flash frozen in liquid nitrogen and kept at −80°C until ready to use. An aliquot of embryos was taken from each batch before freezing to verify staging. For this, embryos were fixed in heptane and 2% paraformaldehyde for 20 minutes with shaking, devitellinized in methanol, washed with methanol and rehydrated in PBS + 0.1% Triton X-100 overnight. Embryos were stained with DAPI and mounted in Vectashield medium (Vector Labs). Images were acquired on a Nikon Ti-E inverted microscope with a Zyla sCMOS digital camera.

### Genomic DNA purification and RNase treatment

Genomic DNA purification and DRIP protocols are based on Alecki and Francis et al. 2020 and Xu and Sun et al. 2017. For genomic DNA isolation from S2 cells, cells were collected at 70-80% confluency, washed once in PBS, resuspended in TE with 0.5% SDS and 100 μg/mL proteinase K and incubated at 37°C overnight. Embryos were devitellinized in heptane and methanol, rinsed thoroughly in PBS and incubated in 50 mM Tris-HCl pH 8.0, 100 mM EDTA, 100 mM NaCl, 0.5% SDS, and 5 mg/ml proteinase K for 3 hours at 50°C. At this point, cells and embryos were processed the same. Extracts were purified with phenol:chloroform, and DNA was precipitated with sodium acetate and ethanol. DNA was spooled using a glass pipette and transferred to 70% ethanol. After several washes in ethanol, the DNA was air dried and resuspended in TE. To degrade free RNA, samples were incubated with 100 μg of RNase A with 500mM NaCl for 1 hour at 37°C. RNase A was degraded by spiking in 100 μg/mL proteinase K and incubated for an additional 45 minutes. Samples were cleaned with phenol:chloroform, precipitated with sodium acetate and ethanol, and resuspended in TE. Samples were diluted to 100 ng/μL and sonicated in a Bioruptor Plus for 8 cycles (30“ on/90” off) on low power. 10 μg of nucleic acid was digested with 5 μL RNase H (NEB) at 37°C for 16 hours and 10 μg was mock digested without RNase H. Both samples had 1 μL of RNase III added (Thermo Fisher). After phenol:chloroform purification and precipitation, samples were immediately used for DRIP or slot blot experiments.

### Slot blot

Hybond Nylon membrane (Amersham) was pre-soaked in TE and a slot blot apparatus was assembled according to manufacturer’s instructions (Bio-Rad). Samples with matching RNase H-digested controls were added to the blot, and nucleic acids were crosslinked to the membrane with a Strategene UV Stratalinker 1800 using the auto crosslink setting. Blots were blocked in milk, incubated with S9.6 (1:2,000) followed by mouse-HRP and imaged in a Bio-Rad Chemidoc MP. After imaging the R-loops, blots were stripped and re-probed using a dsDNA-specific antibody (Abcam ab27156) at 1:20,000. Intensities were measured with ImageJ (Schneider and Eliceiri et al. 2012), and normalized intensity was obtained by dividing the S9.6 signal by the dsDNA signal (Ramirez and Grunseich et al. 2021). Each sample was the average of four technical replicates.

### DRIP-qPCR and ssDRIP-seq

DRIP was carried out as described in Ginno and Chédin et al. 2012. Briefly, 4.4 μg of DNA was resuspended in 500 μL of TE. 10% was taken for the input sample. DRIP binding buffer was added to each sample (10mM sodium phosphate, 140mM NaCl, 0.05% Triton X-100 final concentration) and 20 μL of 1 mg/mL S9.6 was added to each DRIP reaction. After overnight incubation at 4°C, 50 μL of pre-washed protein G Dynabeads (Life Technologies) were added to the extract. After 2 hours at 4°C, beads with captured nucleic acid were washed in 1x DRIP binding buffer 5 times and eluted in 50mM Tris, 10mM EDTA, 0.5% SDS with proteinase K at 50°C for 45 minutes. Nucleic acid in the eluate was purified with phenol:chloroform, precipitated and resuspended in 10mM Tris. For DRIP-qPCR, samples were diluted 1:10 in water and mixed with iTaq (Bio-Rad), with analysis carried out on a Bio-Rad CFX96 Touch instrument. For ssDRIP, nucleic acid was sonicated in a Bioruptor Plus for 8 cycles at high power (30” on/30” off) to 250 bp. Libraries were constructed with the Accel-NGS 1S Plus DNA Library Kit according to the manufacturer’s instruction (Swift Biosciences 10024). Barcoded libraries were sequenced using an Illumina Novaseq for 150bp PE reads.

#### Bioinformatics

##### Alignment and peak calling

Fastq files were initially trimmed of adapters using Trimmomatic v0.3.8 (Bolger and Usadel et al. 2014). Each paired read was trimmed 10 base-pairs at the 3’ end to eliminate the additional low complexity from the library preparation kit. Reads for sequencing were mapped to the Drosophila genome (dm6) using bowtie2 version 2.3.4.1 using the –very-sensitive-local setting (Langmead and Salzberg 2012). Duplicates were marked using picard MarkDuplicates v2.17.10, and stranded bam files were created using samtools as described in Xu and Sun et al. 2017 (Li and Durbin et al. 2009). Stranded bam files were used to generate ssDRIP peaks with callpeaks from MAC2 v2.1.2 (Zhang and Liu et al. 2008). The RNase H pretreated DRIP file was used as control, peak calling was done in paired-end mode, with –keep-dup=auto and effective genome size for Drosophila dm6. Stranded reads were visualized using deeptools bamCoverage using --binSize 50bp, --ignoreForNormalization chrY chrM, and --normalizeUsing RPKM (Ramírez and Manke et al. 2014). A small number of reads mapped to both strands. These reads were discarded for the analysis.

##### ssDRIP-seq analysis

Annotation of R-loop peaks was done with HOMER software package using annotatePeaks.pl (Heinz and Glass 2010). Stranded R-loops were determined via bedtools intersect with strandedness against the Refseq Drosophila transcriptome, downloaded from UCSC genome browser. Metagene plots were made with the Deeptools software package, using computeMatrix and plotProfile. GRO-seq FPKM counts were determined with HOMER analyzeRepeats.pl using S2 datasets from Core and Lis et al. 2012 and GRO-seq data on 2-2.5 embryos from Saunders and Ashe et al. 2013.

##### Functional genomic data from modENCODE

We downloaded histone modification peaks and transcription factor binding sites identified by ChIP-chip or ChIP-seq in Drosophila from ModENCODE (Contrino and Hu et al. 2012). We considered samples assayed in S2 cells and at two developmental timepoints (2-4hr, 14-16hr). These were chosen to match the ssDRIP timepoints.

**Table 1.**
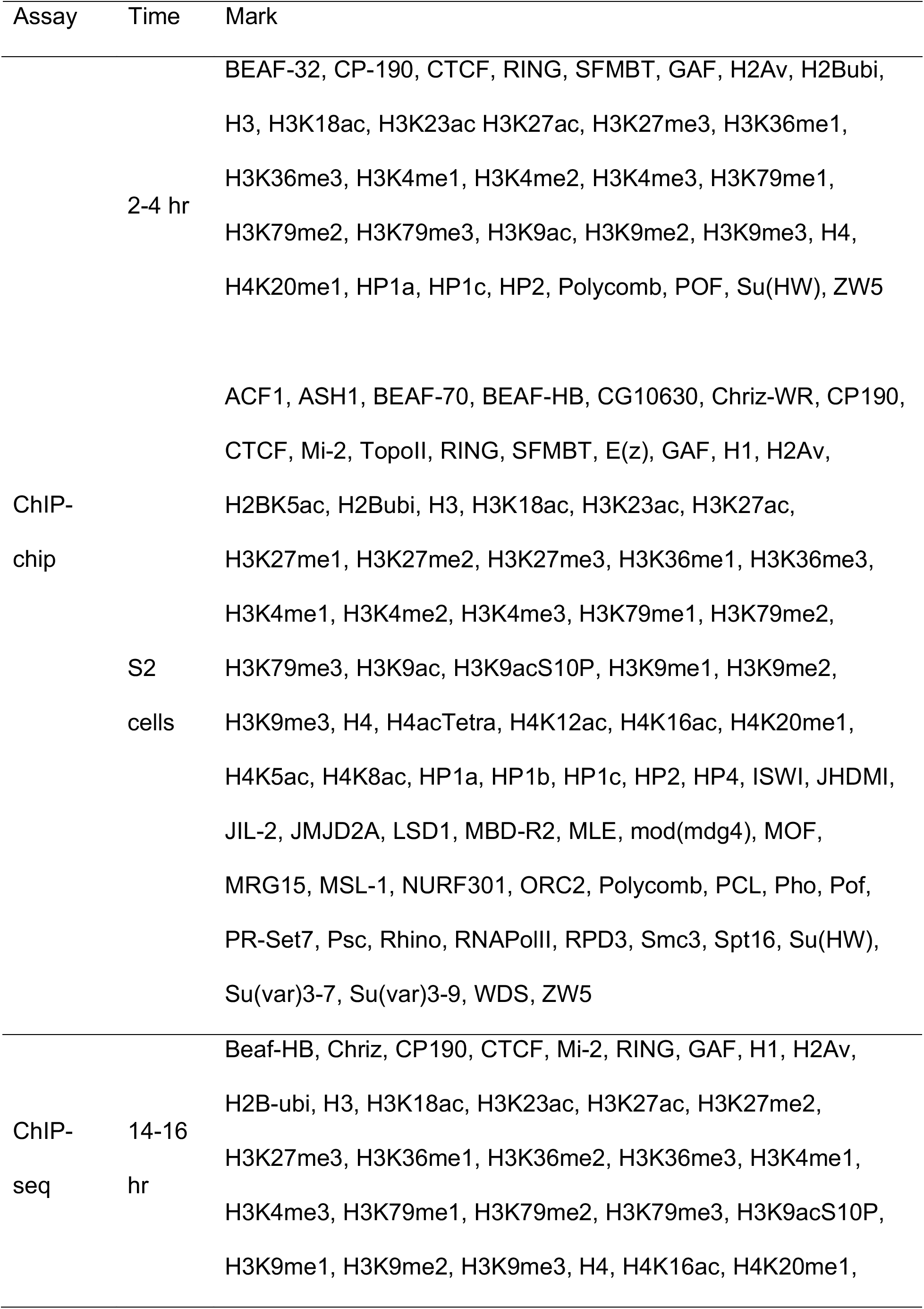
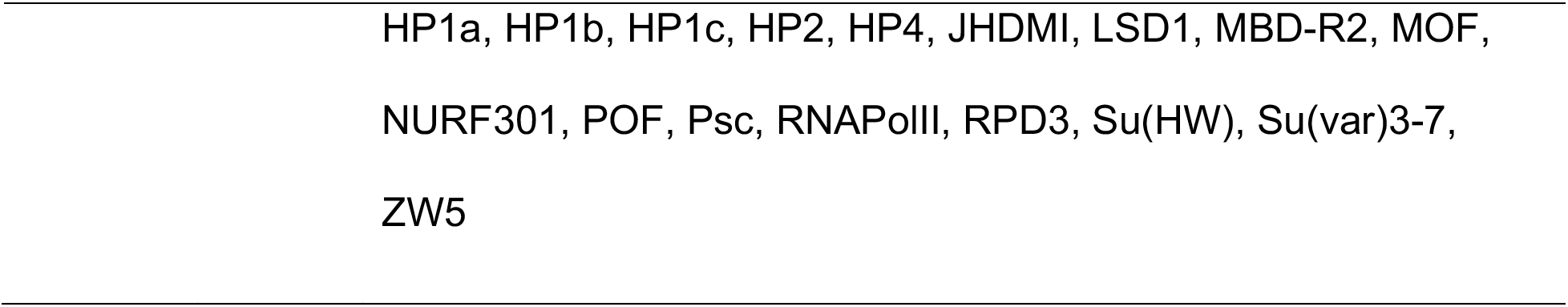
List of available ChIP-chip and ChIP-seq from modENCODE.

##### Chromatin marker enrichment in R-loops

For each ChIP-chip or ChIP-seq marker with a matching DRIP timepoint, we calculated the number of overlapping base-pairs (bp) between the marker and the R-loop peaks. We used permutation-based approach to determine whether the observed amount of overlap was more or less than expected by chance. Briefly, we calculated an empirical *p* value for the observed amount of overlap by comparing the number of overlapping bp to a null distribution. We obtained the null distribution by randomly shuffling length-matched regions throughout the genome and calculating the amount of overlap in each permutation. The *p*-values are adjusted for multiple testing using the Bonferroni method.

When permuting, we matched the length distribution of the shuffled peaks to the original set of peaks, and excluded all gap and blacklisted regions from consideration (dm3; version 1) (Amemiya and Boyle et al. 2019). Peaks called from DRIP were lifted over to dm3 for this analysis. For peaks obtained from ChIP-chip data, we required that the shuffled peaks maintained both the overall length distribution and the probe density of the original peak. We reshuffled any peaks that fell more than 2 standard deviations (approx. 0.03) away from the original probe density until at least 99% of the original peaks were appropriately matched. We performed 1000 permutations for each marker and R-loop pair.

For the general analyses, we maintained the location of the R-loop peaks and shuffled the locations of the histone modification or transcription factor binding peaks. For a secondary analysis, we examined a subset of R-loops quantified specifically in the TTS and 3’ UTR. For this set of R-loops, we maintained the R-loop location within the TTS/3’ UTR and shuffled the chromatin markers.

##### Calculation of GC-skew in R-loops

We calculated GC skew over three sets of genomic regions: (1) all of the ascertained R-loops, (2) all genes that do not overlap R-loops, and (3) all genes that overlap R-loops. We used the bedTools suite to obtain sequences for each of these genomic regions before calculating skew (Quinlan and Hall 2010). GC skew was calculated for 50 bp windows tiled across the annotation regions as 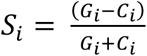 (McLean and Devine et al. 1998).

In the equation, *G_i_* represents the frequency of guanine nucleotides and *C_i_* represents the frequency of cytosine nucleotides in the window *i*. The range of GC skew for a window (*S_i_*) spans from −1 to 1. The resulting GC skew across each set of genomic regions was plotted using deepTools.

## DATA ACCESS

Data sets generated in this study can be found under the GEO accession number: GSE185403.

## ACKNOWLEDGEMENTS

We thank the Vanderbilt VANTAGE core for Illumina sequencing and the Vanderbilt Antibody and Protein Resource core for purifying the S9.6 antibody. The Vanderbilt Antibody and Protein Resource core is supported by the Vanderbilt Institute of Chemical Biology and the Vanderbilt Ingram Cancer Center (P30 CA68485). We thank Martina Brienza-Ramos for cloning of the RNase H1 plasmids used for fly injections. We thank Emily Hodges, Robin Armstrong and Frederic Chédin for providing critical feedback on the manuscript. We thank Lionel Sanz, Célia Alecki and Nicole Francis for technical advice. This work was supported by National Institutes of Health (NIH) General Medical Sciences awards R35GM127087 to JAC and R35GM128650 to JTN.

## AUTHOR CONTRIBUTIONS

AM and JTN planned and designed the research; AM performed experiments; AM and MB analyzed data with supervision from JAC; AM and JTN wrote the manuscript. AM, MB, JAC and JTN edited the manuscript.

