## Supplemental Material for "R-loop mapping and characterization during Drosophila embryogenesis reveals developmental plasticity in R-loop signatures"

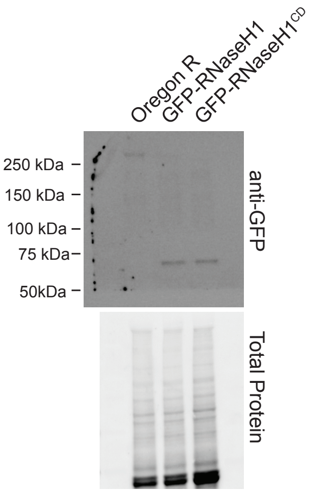


Supplemental Figure 1: Expression of RNaseH1 constructs in early embryos. Western blot (anti-GFP) showing maternally deposited GFP-RNaseH1 and GFP-RNaseH1$^{CD}$ in 0-6 hour embryos. Expected size of GFP + RNaseH1 = 65.1 kDa.

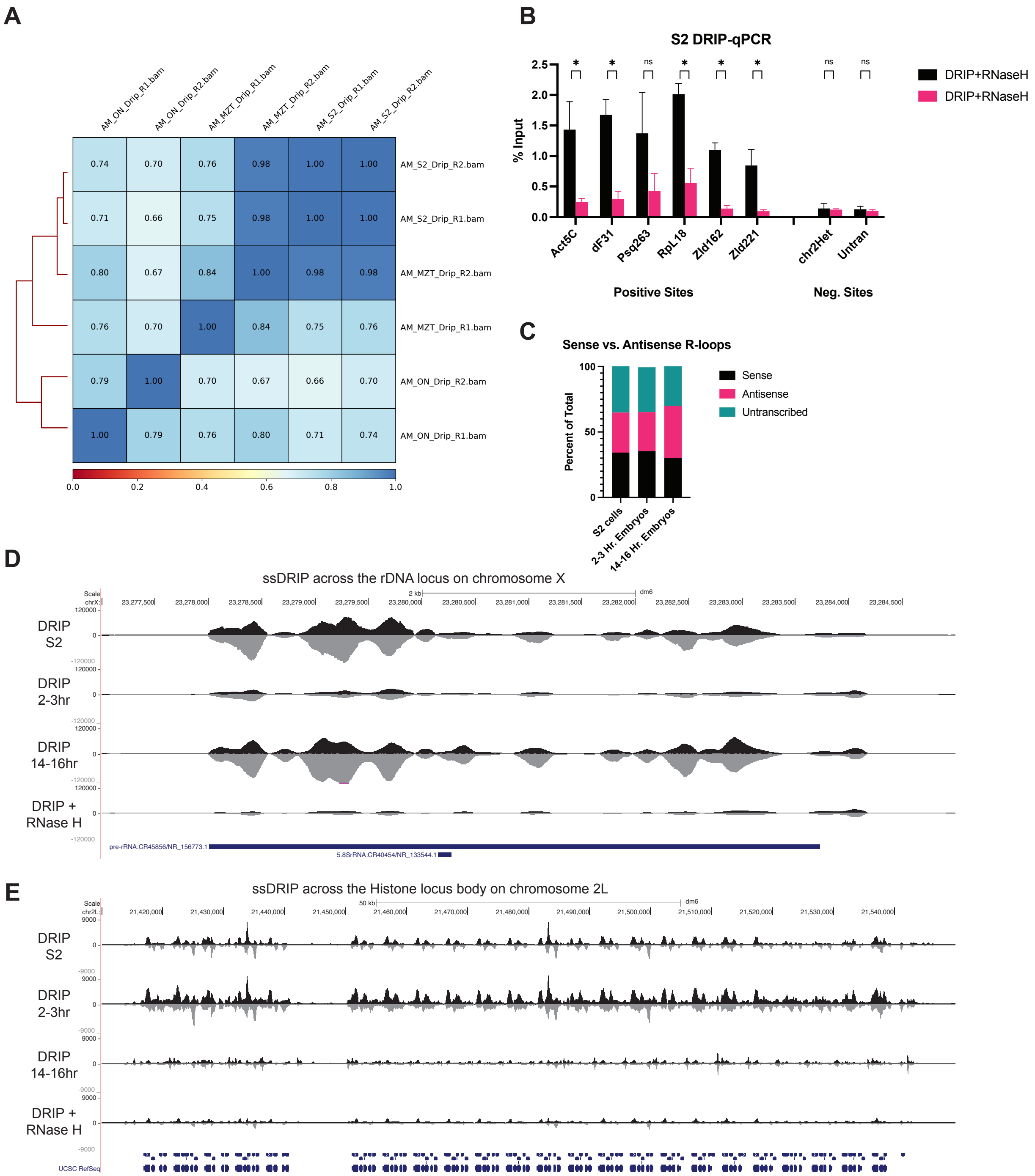


Supplemental Figure 2: Properties of R-loops in Drosophila (A) Correlation between ssDRIP-seq replicates at 1kb resolution. (B) DRIP-qPCR validation of several R-llop positive and negative loci in S2 cells. (C) Quantification of the percent of R-loops mapping to sense, antisense and unannotated regions of the genome. (D) R-loop abundance at a rRNA locus (note the difference in scale compared to profiles in Fig. 2). (E) R-loop abundance at the histone locus on chromosome *2L.*

**A**

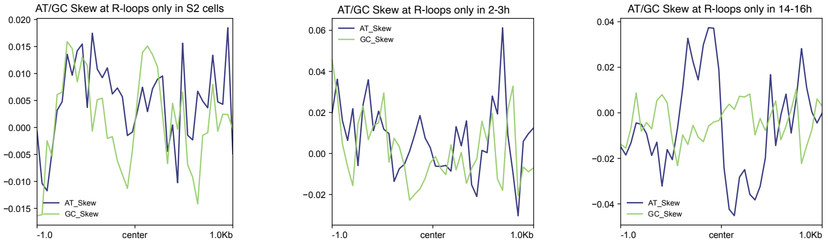


**B**

| Rank | Motif | P-value | log P-pvalue | % of Targets | % of Background | STD(Bg STD) | Best Match/Details |
|---|---|---|---|---|---|---|---|
| 1 | 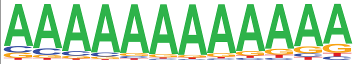 | 1e-1450 | -3.340e+03 | 21.44% | 8.73% | 48.4bp (78.7bp) | hb/dmmpmm(Noyes)/fly(0.794) More Information \| Similar Motifs Found |
| 2 | 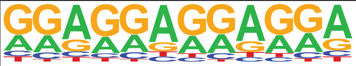 | 1e-961 | -2.215e+03 | 31.82% | 18.65% | 51.4bp (70.0bp) | Trl/dmmpmm(Down)/fly(0.559) More Information \| Similar Motifs Found |
| 3 | 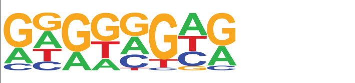 | 1e-149 | -3.432e+02 | 42.12% | 36.14% | 56.1bp (64.8bp) | Trl(Zf)/S2-GAGAfactor -ChIP-Seq(GSE40646)/Homer(0.647) More Information \| Similar Motifs Found |
| 4 | 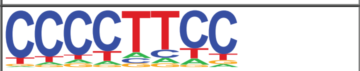 | 1e-127 | -2.947e+02 | 27.92% | 23.00% | 55.2bp (68.4bp) | Kr/dmmpmm(Noyes)/fly(0.718) More Information \| Similar Motifs Found |
| 5 | 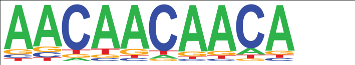 | 1e-106 | -2.457e+02 | 14.88% | 11.43% | 56.1bp (66.6bp) | Aef1/dmmpmm(Pollard)/fly(0.864) More Information \| Similar Motifs Found |
| 6 | 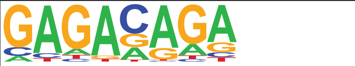 | 1e-104 | -2.398e+02 | 40.18% | 35.22% | 56.3bp (67.1bp) | Trl/MA0205.2/Jaspar(0.781) More Information \| Similar Motifs Found |
| 7 | 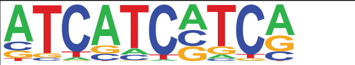 | 1e-53 | -1.239e+02 | 7.78% | 5.96% | 56.7bp (57.1bp) | ttk/MA0460.1/Jaspar(0.615) More Information \| Similar Motifs Found |
| 8 | 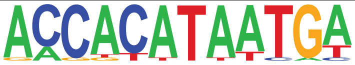 | 1e-52 | -1.201e+02 | 0.13% | 0.01% | 58.2bp (42.2bp) | Btn/dmmpmm(Noyes_hd)/fly(0.690) More Information \| Similar Motifs Found |
| 9 | 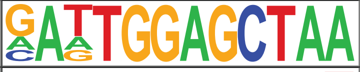 | 1e-38 | -8.804e+01 | 0.08% | 0.00% | 53.6bp (31.1bp) | POL013.1_MED-1/Jaspar(0.622) More Information \| Similar Motifs Found |
| 10 | 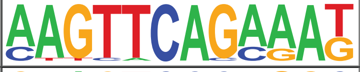 | 1e-37 | -8.600e+01 | 0.12% | 0.01% | 55.1bp (27.4bp) | ct/MA0218.1/Jaspar(0.627) More Information \| Similar Motifs Found |
| 11 | 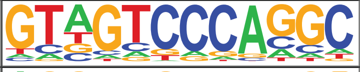 | 1e-36 | -8.456e+01 | 0.07% | 0.00% | 51.8bp (18.8bp) | dl-B/dmmpmm(Bergman)/fly(0.559) More Information \| Similar Motifs Found |
| 12 | 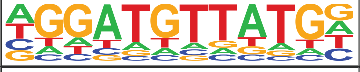 | 1e-36 | -8.456e+01 | 0.07% | 0.00% | 53.5bp (38.4bp) | ara/dmmpmm(Noyes_hd)/fly(0.712) More Information \| Similar Motifs Found |
| 13 | 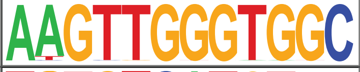 | 1e-34 | -7.985e+01 | 0.09% | 0.01% | 52.3bp (18.5bp) | Hr46/dmmpmm(Pollard)/fly(0.574) More Information \| Similar Motifs Found |
| 14 | 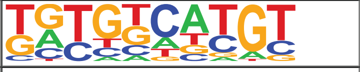 | 1e-34 | -7.935e+01 | 2.89% | 2.02% | 56.0bp (60.3bp) | h/dmmpmm(Noyes)/fly(0.670) More Information \| Similar Motifs Found |
| 15 | 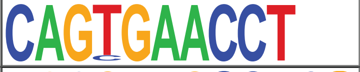 | 1e-33 | -7.768e+01 | 0.07% | 0.00% | 56.8bp (15.9bp) | eyg/dmmpmm(Bergman)/fly(0.689) More Information \| Similar Motifs Found |
| 16 | 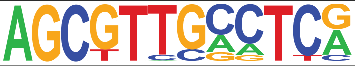 | 1e-30 | -7.018e+01 | 0.09% | 0.01% | 55.7bp (14.8bp) | POL010.1_DCE_S_III/Jaspar(0.589) More Information \| Similar Motifs Found |

Supplemental Figure 3: Sequence properties of R-loops in Drosophila (A) Metaplots of AT and GC skew at developmental-specific sites of R-loop formation (note the difference in scale for each window). (B) The top 16 results from the HOMER motif analysis. Only P-values less than 1e-100 should be considered as potential motifs.
